# Myonuclear loss, rather than senescent myonuclei, associates with fiber type-specific atrophy in aging human skeletal muscle

**DOI:** 10.64898/2026.02.11.705446

**Authors:** Carlos S. Zepeda, Isabell Dobrzycki, Paige N. Acklie, Cory M. Dungan, Ronald G. Jones, Kevin A. Murach, Christopher W. Sundberg

**Author notes:** Address for Correspondence: Christopher W. Sundberg, Ph.D., University of Wisconsin – Madison, Wisconsin Institutes for Medical Research, Room 3405, 1111 Highland Avenue, Madison, WI 53705-2275.

## Abstract

Age-related reductions in whole-muscle function are attributed, in part, to pronounced atrophy of muscle fibers expressing the fast myosin heavy chain (MyHC) II isoforms. Senescence, a state of irreversible cell cycle arrest that can be characterized by DNA damage (γH2AX) and chromatin remodeling (loss of nuclear HMGB1), may contribute to skeletal muscle aging. Muscle nuclei (myonuclei) maintain fiber size and function and could exhibit senescence-associated features; however, the prevalence of senescent myonuclei and whether they contribute to fast fiber atrophy in older adults remains unknown. *Vastus lateralis* biopsies from 20 young (19-34yr; 10 females) and 20 older (65-84yr; 10 females) adults were analyzed via immunohistochemistry for myonuclei positive for γH2AX (γH2AX^+^) and negative for HMGB1 (HMGB1^−^). MyHC II cross-sectional area (CSA) was ∼70% larger in young compared with old, whereas MyHC I CSA did not differ with age. The relative abundance of γH2AX^+^/HMGB1^−^ myonuclei did not differ with age and was not associated with CSA in either fiber type. Single-nucleus RNA-sequencing corroborated no age-related difference in the prevalence of myonuclei with senescence-associated features. Myonuclear content of MyHC II fibers was ∼30% higher in young compared with old and was closely associated with CSA in both fiber types. Size-cluster analysis revealed a pronounced age-related leftward shift in MyHC II CSA that paralleled the reductions in myonuclear number, consistent with myonuclear loss. These data suggest that age-related fast fiber atrophy is not attributed to an increased prevalence of senescent myonuclei but instead occurs concomitantly with fiber type-specific myonuclear loss across the lifespan.

## INTRODUCTION

Skeletal muscle constitutes ∼40% of total body mass and plays an essential role in locomotion, maintaining posture, breathing, and regulating metabolism, all of which are essential for maintaining an active and independent lifestyle. Importantly, beginning around the fourth decade of life, skeletal muscle mass declines at a rate of ∼0.3–0.6 % /yr (Mitchell et al., 2012; Naruse et al., 2023). This progressive age-related loss of muscle mass is clinically significant because it can contribute to declines in mobility, physical function, and independence. A major factor contributing to the age-related loss in muscle mass is the atrophy of muscle fibers, particularly in fibers expressing the fast myosin heavy chain (MyHC) II isoforms, which are more susceptible to age-related reductions in size and function than slow MyHC I fibers (Grosicki et al., 2022; Lexell et al., 1988; Nilwik et al., 2013; Sundberg et al., 2024; Teigen et al., 2020). Previous studies report that fast MyHC II fiber atrophy is a major contributor to the pronounced age-related reduction in fast fiber force and power, which has important implications for whole muscle function (Gries et al., 2019; Grosicki et al., 2021; Grosicki et al., 2022; Sundberg, Hunter, et al., 2018; Sundberg et al., 2024; Teigen et al., 2026). Despite the importance of preserving MyHC II fiber size and overall muscle mass with advancing age, the mechanisms underlying age-related muscle loss are not well understood.

Emerging evidence suggests that cellular senescence, a stress-induced response caused by DNA damage and chromatin remodeling that results in irreversible cell cycle arrest and a deleterious senescent associated secretory phenotype, or “SASP”, may contribute to the loss of muscle mass and function with aging (Dungan et al., 2023; Englund et al., 2021). Senescent cells are resistant to apoptosis and tend to persist in a tissue, often causing widespread local and systemic dysfunction (Ogrodnik et al., 2019). Skeletal muscle is unique in that muscle fibers are large, multinucleated cells, and myonuclei are non-dividing organelles important for maintaining fiber structure and function. Thus, hallmark features of cellular senescence, such as persistent DNA damage and altered chromatin structure, may compromise myonuclear DNA integrity and contribute to age-related fiber atrophy. In myonuclei, persistent DNA damage can be visualized by the presence of γH2AX, the phosphorylated form of the histone H2A variant that signals double-stranded break sites and facilitates the recruitment and assembly of DNA repair machinery. The sustained presence of γH2AX foci is therefore widely used as an indicator of senescent cells (Bernadotte et al., 2016). Another senescence-associated marker related to the nucleus is the nuclear-to-cytoplasmic translocation of high mobility group B1 (HMGB1), a chromatin-associated protein that binds to DNA and histones to regulate chromatin structure and transcription factor accessibility (Tang et al., 2023). When HMGB1 exits the nucleus, it serves as a damage-associated molecular pattern that promotes an inflammatory response (Bianchi, 2009). Previous research has shown that HMGB1 is translocated out of the nucleus following a senescence-inducing stimulus, and therefore, the loss of nuclear HMGB1 (HMGB1^−^) is also used as a marker of cellular senescence (Davalos et al., 2013). Together, characterizing the localization of γH2AX and HMGB1 within skeletal muscle myonuclei may provide insight into whether compromised myonuclear DNA integrity and cellular senescence is contributing to age-related muscle fiber atrophy. Furthermore, using multiple markers for identifying senescent features aligns with current recommendations and ensures a more rigorous and robust analysis (Gonzalez-Gualda et al., 2021; Gorgoulis et al., 2019; Ogrodnik et al., 2024).

Only a few studies have investigated the presence of γH2AX or HMGB1 in skeletal muscle, and the findings are inconsistent. Zhang et al. (2022) reported a greater abundance of HMGB1^−^nuclei in older mice, along with more telomere-associated foci in both older mice as well as older humans when compared with younger cohorts. Interestingly, they also found a higher expression of other genes associated with senescence (*p16* and *p21*) in older mice within muscles composed primarily of fast fibers, such as the tibialis anterior and extensor digitorum longus (Zhang et al., 2022). These preclinical findings suggest a muscle- and/or fiber type-specific vulnerability to senescence and are consistent with earlier work showing that an increased prevalence of HMGB1^−^nuclei in older mice was associated with reduced muscle fiber size (da Silva et al., 2019). In contrast, Dungan et al. (2020) reported no age-related differences in the prevalence of γH2AX-positive (γH2AX^+^) myonuclei in skeletal muscle from young and older adults (Dungan et al., 2020). Although these studies provide valuable insight into the effects of cellular senescence within skeletal muscle, there are methodological limitations that should be considered when interpreting the findings: 1) a muscle morphology marker was not included to delineate fiber borders and distinguish myonuclei from other mononuclear cells (da Silva et al., 2019; Zhang et al., 2022), and 2) fiber types were not analyzed separately (Dungan et al., 2020). To our knowledge, no studies have investigated whether older adults exhibit an increased prevalence of myonuclei expressing multiple markers of cellular senescence in a fiber type-specific manner. Thus, the primary aim of this study was to test the hypotheses that the MyHC II fibers from older adults would have an increased prevalence of γH2AX^+^/HMGB1^−^ myonuclei and that this would be associated with their smaller size. To support the histology data on our large cohort, we also analyzed single-nucleus RNA-sequencing (snRNA-seq) data obtained from skeletal muscle of young and older males (Perez et al., 2022), and examined whether the expression of senescence-associated markers differed with age.

In addition to the presence of senescent myonuclei, changes in myonuclear number may have important implications for fiber type-specific atrophy with aging. Myonuclei feature marked transcriptional heterogeneity (Petrany et al., 2020) and play critical roles in mediating exercise adaptations (Koopmans et al., 2022) and regulating cell growth (Borowik et al., 2024); thus, the loss of myonuclei with atrophy and/or aging could have consequences for skeletal muscle size and function. Indeed, several studies have demonstrated associations between myonuclear content and both slow and fast fiber size (Horwath, Envall, et al., 2021; Horwath, Moberg, et al., 2021; Snijders et al., 2020; Verdijk et al., 2010). However, whether fiber atrophy is accompanied by a reduction in myonuclear content remains unresolved and has been extensively debated (Kirby & Dupont-Versteegden, 2022; Kirby & Dupont - Versteegden, 2022; L. Schwartz & K. Gundersen, 2022; L. M. Schwartz & K. Gundersen, 2022). Given our large histological dataset across fiber types, a secondary aim was to determine whether age-related fast fiber atrophy in older males and females is accompanied by a reduction in myonuclear content. We hypothesized that fast fibers from older adults would exhibit reduced myonuclear number, with no age differences in slow fibers, and that myonuclear number would be positively associated with fiber size irrespective of biological sex.

## METHODS

### Participants and Ethical Approval

Twenty young (19-34 yr; 10 females/10 males) and 20 older adults (65-84 yr; 10 females/10 males) volunteered and provided written informed consent to participate in this study. Participants underwent a general health screening that included an assessment of body composition and thigh lean mass via dual X-ray absorptiometry (Lunar iDXA; GE, Madison, WI) (Sundberg, Kuplic, et al., 2018). Participants were healthy, community-dwelling adults free of any known neurological, musculoskeletal, and cardiovascular diseases and were excluded from participation if they had any known major health concerns. All experimental procedures were approved by the Marquette University Institutional Review Board and conformed to the principles in the Declaration of Helsinki. Physical activity levels and anthropometric data are reported in Table 1.

**Table 1:** Participant anthropometrics and physical activity levels.

| Variable | Units | Males |  | Females |  | P-value |  |  |
| --- | --- | --- | --- | --- | --- | --- | --- | --- |
|  |  | Young (10) | Old (10) | Young (10) | Old (10) | Age | Sex | Age*Sex |
| Age | yr | 27.0 ± 5.8 | 74.1 ± 6.6 | 23.0 ± 2.7 | 70.6 ± 4.2 | **<0.001** | **0.025** | 0.859 |
| Height | cm | 179.6 ± 7.1 | 176.3 ± 8.0 | 162.3 ± 8.5 | 161.0 ± 6.7 | 0.348 | **<0.001** | 0.702 |
| Weight | kg | 81.3 ± 10.9 | 82.7 ± 5.5 | 64.7 ± 9.9 | 65.3 ± 12.4 | 0.747 | **<0.001** | 0.890 |
| BMI | kg·m<sup>-2</sup> | 25.2 ± 2.5 | 26.7 ± 2.2 | 24.5 ± 3.1 | 25.0 ± 3.4 | 0.258 | 0.196 | 0.598 |
| Body Fat | % | 22.2 ± 3.5 (9) | 31.6 ± 4.9 | 31.2 ± 8.0 | 36.3 ± 5.8 | **<0.001** | **<0.001** | 0.263 |
| Thigh Lean Mass | kg | 8.9 ± 2.2 (9) | 6.4 ± 1.2 | 5.5 ± 1.0 | 4.7 ± 1.2 | **0.001** | **<0.001** | 0.073 |
| Physical Activity | steps/day | 8842 ± 3446 (9) | 6803 ± 2733 | 7953 ± 3539 | 8559 ± 4955 | 0.556 | 0.721 | 0.281 |

Values are reported as means ± SD. Body fat percentage was measured via dual X-ray absorptiometry (Lunar iDXA, GE, Madison, WI). Physical activity was measured via triaxial accelerometry (GT3X, ActiGraph, Pensacola, FL). The sample sizes (n) for each cohort and certain variables are reported in parentheses. A significant age or sex difference is indicated in bold when $P < 0.05$ .

### Physical activity assessment

Physical activity for each participant was quantified using a triaxial accelerometer (GT3X; ActiGraph, Pensacola, FL, USA) worn around the waist for at least 4 days (2 weekdays and 2 weekend days) as reported previously (Hassanlouei et al., 2017; Sundberg, Hunter, et al., 2018). The data were recorded for each participant if the accelerometer was worn for a minimum of 8 hours on at least 3 days (Hart et al., 2011).

### Muscle biopsy

A muscle biopsy from the *vastus lateralis* was obtained from each participant as described previously (Bergstrom, 1975; Sundberg, Hunter, et al., 2018). Participants were instructed to abstain from strenuous exercise of the lower limbs for 48 h prior to the biopsy and arrive to the laboratory in the morning fasted and without the consumption of caffeine for ≥ 8 h. The biopsy location was cleaned with 70% ethanol, sterilized with 10% povidone-iodine and anaesthetized with 1% lidocaine HCl. A small ∼1 cm incision was made overlying the distal 1/3 of the muscle belly, and the biopsy needle inserted under local suction to obtain the tissue sample. Bundles from the biopsy sample were oriented longitudinally on a small notecard and immediately frozen in liquid nitrogen-cooled isopentane and stored at -80°C until sectioning.

### Immunohistochemistry

Immunohistochemistry (IHC) analysis was performed to determine fiber type-specific cross-sectional area (CSA) and prevalence of myonuclei identified as senescent (γH2AX^+^/HMGB1^−^, DAPI^+^, and within dystrophin). Muscle samples were removed from the -80°C and placed into a cryostat (HM525NX, Thermo Scientific) set at -21°C. After the tissue equilibrated to -21°C (≥30 min), 7 μm sections were cut at a 10° clearance angle and retrieved with Superfrost Plus slides (Ref. 6776214, Epredia). Sections were air-dried at room temperature (RT) for ≥1 hr and then stored at -80°C until staining. For each analysis, slides were taken from -80°C, air dried for ≥ 30 min at RT, and sections circled with an ImmEdge pen (H-4000, Vector Laboratories) to form a hydrophobic barrier which air dried for ≥15 min.

### Fiber type-specific γH2AX and HMGB1immunodetection

This analysis was performed according to a protocol modified from Dungan et al. (2020). Muscle sections were fixed in 4% paraformaldehyde for 5 min followed by 3x3 min 1X PBS washes. Sections were then blocked in 1% Triton X-100 and 2% bovine serum albumin (BSA) diluted in 1X PBS for 60 min. Following blocking, sections were incubated in a 1°Ab cocktail consisting of Ms IgG2a α-dystrophin (1:100; MANDYS1(3B7); DSHB, RRID: AB_528206), Ms IgG2b α-MyHC I (1:100, BA-D5-c; DSHB, RRID: AB_2235587), Ms IgG1 α-γH2AX (1:1000, 05-636, Millipore Sigma, RRID: AB_309864), and Rb IgG α-HMGB1 (1:250, ab18256; Abcam, RRID: AB_4443620) diluted in 1% Triton X-100 and 2% BSA blocking buffer for 90 min at RT. Muscle sections were then washed 3x5 min with fresh 1X PBS and incubated in a 2°Ab cocktail consisting of Gt α-Ms IgG2a AF488 (1:250, A21131, Invitrogen, RRID: AB_2535771), Gt α-Ms IgG2b AF488 (1:250, A21141, Invitrogen, RRID: AB_2535778), Gt α-Ms IgG1 AF555 (1:250, A21127; Invitrogen, RRID: AB_2535769), and Gt α-Rb IgG AF647 (1:250, A21245; Invitrogen, RRID: AB_2535813) diluted in 1% Triton X-100 and 2% BSA blocking buffer for 60 min at RT. Sections were then washed 3x5 min in 1X PBS and incubated for 10 min in DAPI (1:10,000, D1306; Invitrogen). Finally, sections were washed 3x5 min in 1X PBS and mounted with covers slips using 1:1 1X PBS/glycerol. Slides were stored at 4 °C until imaging.

### Image acquisition and analysis

Images were acquired using a high-resolution fluorescence microscope (BZ-X810, Keyence) with a Plan Apochromat 20x mounted objective (BZ-PA20, Keyence) and filters DAPI (49000, Chroma Technology), Cy5 (49006, Chroma Technology), FITC/Alexa Fluor 488/Fluo3/Oregon Green (49011, Chroma Technology), and CY3/TRITC (49004, Chroma Technology). Regions of the cross-section that appeared damaged or distorted were manually excluded from all analyses.

From the γH2AX/HMGB1 staining, myonuclear content was determined for each fiber type (MyHC I and MyHC II) by a trained investigator. Myonuclei were identified as having >50% of the nucleus area residing within the dystrophin border (Bruusgaard & Gundersen, 2008). Centrally located nuclei (i.e., centralized nuclei) fully enclosed within the dystrophin border but not located at the periphery of the fiber were not included in the analysis of myonuclear content (Snijders et al., 2021). Myonuclear domain was calculated as fiber CSA divided by the number of myonuclei of that fiber. The myonuclear analysis included a minimum of 100 fibers of each type per cross-section, or all available fibers when <100 of a given fiber type were present. Accordingly, 4 participants had fewer than 100 MyHC I fibers available for analysis (66–90 fibers) and 9 participants had fewer than 100 MyHC II fibers (33–93 fibers). In total, 4,218 MyHC I fibers (Young = 2,005; Old = 2,213) and 4,055 MyHC II fibers (Young = 2,022; Old = 2,033) were included in the myonuclear analysis. Fibers were further stratified into 500 μm^2^ size bins for size-cluster analyses, with fiber counts pooled across males and females within each age group (Table A1). Myonuclei positive for γH2AX (γH2AX^+^) and negative for HMGB1 (HMGB1^−^) were considered senescent (γH2AX^+^/HMGB1^−^) (Fig. 1A-C). γH2AX^+^/HMGB1^−^ myonuclei were manually counted using ImageJ (National Institute of Health, USA).

**Fig. 1.**
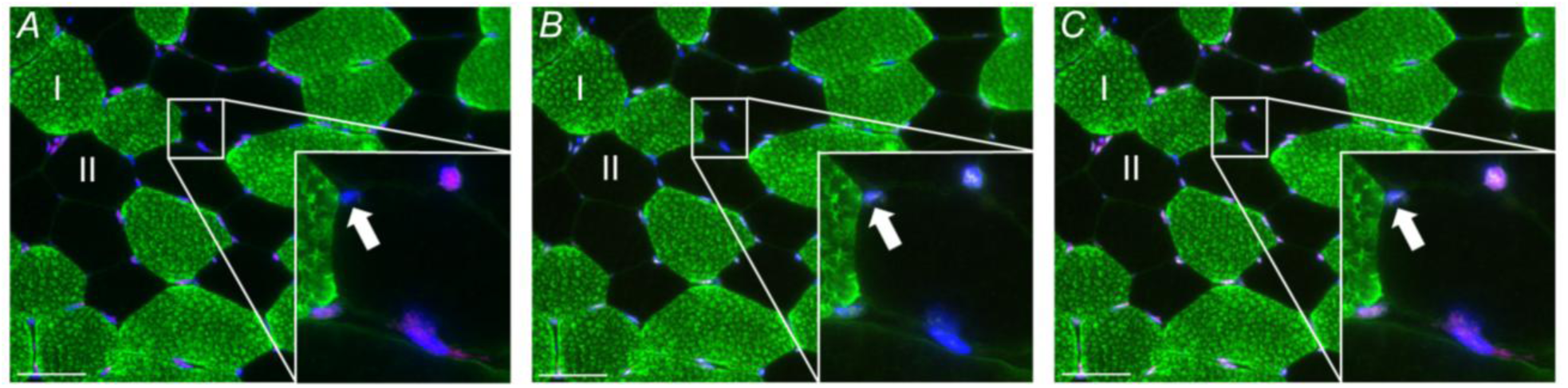
Representative images of (A) HMGB1 (pink), (B) γH2AX (white), and (C) γH2AX/HMGB1 overlayed. The insets in panels A-C show a magnified region of each image with the white arrow indicating a myonucleus that is γH2AX^+^/HMGB1^−^. Myonuclei were identified as senescent if >50% of the nucleus area was within the dystrophin border (green), negative for HMGB1 and positive for γH2AX (γH2AX^+^/HMGB1^−^). Muscle fibers negative for MyHC I (green) were identified as MyHC II. Scale bar is 50 μm.

### Single-nucleus RNA-sequencing

Previously published (Perez et al., 2022) single-nucleus RNA-sequencing of human vastus lateralis muscle biopsies from young and older males were reanalyzed using the Cell Ranger filtered feature barcode matrices provided by the original authors (GSE167186), which were mapped to the human GRCh38 reference genome. Downstream analyses were performed in R (version 4.5.0) using Seurat (version 5.0.0) (Hao et al., 2024). For each sample, nuclei expressing fewer than 200 detected genes were removed, consistent with the low-representation filtering criteria described in the original publication. Additional nucleus-level quality control was performed by excluding nuclei with low or extreme transcript counts or >5% of transcripts mapping to mitochondrial genes. Genes that were expressed in fewer than 3 nuclei were also removed (likely ambient RNA). Putative doublets were identified on the pooled dataset using scDblFinder (version 1.22.0) with an expected doublet rate of 0.075 (Germain et al., 2021), and nuclei labeled as doublets were removed prior to normalization and clustering. Differences in the number of nuclei retained relative to the original report reflect the application of nucleus-level filtering performed to ensure quality of nuclei in our reanalysis of the data.

Normalized expression values and variable features were computed using SCTransform as implemented in Seurat, followed by principal component analysis (PCA). Integration across samples was performed using Harmony (version 1.2.4) with sample identity specified as the integration variable to account for high sample-to-sample variability prior to clustering (Korsunsky et al., 2019). Unsupervised clustering was performed using the Leiden algorithm (leiden version 0.4.3.1) applied to a shared-nearest-neighbor (SNN) graph constructed from the Harmony-corrected embedding. To determine the optimal number of clusters, a resolution sweep (0.2–1.2) was performed and the final resolution (0.2) was selected based on quantitative clustering diagnostics, including graph modularity and average silhouette width. Both the resolution value and number of clusters (ndims = 10) were concordant with the values described in the original study. Cluster identities derived from Leiden were retained and subsequently annotated using established marker genes for skeletal muscle-associated nuclear populations. Differential expression between young and old groups was performed using Wilcoxon rank-sum testing as implemented in Seurat’s FindMarkers() function, consistent with the original study’s approach to age-associated differential expression within nucleus types. Statistical significance was assessed following multiple testing correction using an adjusted *P* value threshold of 0.05.

### Statistical analysis

Anthropometrics, physical activity levels, and fiber type-specific measurements were compared between age groups (young or old) and sex (males or females) with a two-way univariate ANOVA. For analyses correlating the relative abundance of γH2AX^+^/HMGB1^−^ myonuclei and overall myonuclear content with fiber CSA, Pearson correlations were performed. Normal distributions and homogeneity of variance were assessed before any statistical comparisons using the Shapiro-Wilks test and Levene’s statistic, respectively. When necessary, data were transformed to meet assumptions of normality and homogeneity of variance. For the size-cluster analyses, univariate analyses of covariance (ANCOVA) were performed using mean myonuclear content or domain from each cluster, with mean fiber size of each cluster as a covariate, to determine whether the relationship with fiber size differed between young and old. Analyses were performed within each fiber type and the age x fiber size interaction is reported in the text. Only size clusters that contained a minimum of 10 fibers for both young and old were included in the statistical analysis. Based on this criterion, the 1500-2000 μm^2^ to 8500-9000 μm^2^ size clusters were included for MyHC I fibers and the 1000-1500 μm^2^ to 7000-7500 μm^2^ size clusters for MyHC II fibers (Table A1). Statistical analyses were performed using SPSS version 29.0 (IBM Corp., Armonk, NY, USA). Statistical significance was set at *P* < 0.05. Data are presented as mean ± SD in the text and tables and means ± SEM in the figures. Results comparing the two age groups have males and females combined and results comparing the sexes have the young and old combined.

## RESULTS

### Cross-Sectional Area

Cross-sectional area (CSA) for the MyHC I fibers did not differ between young (4679 ± 1182 μm^2^) and old (4524 ± 1002 μm^2^; *P* = 0.635), but males (5000 ± 1023 μm^2^) had ∼19% larger MyHC I fibers compared with females (4203 ± 1016 μm^2^; *P* = 0.019) (Fig. 2A). CSA for the MyHC II fibers was ∼70% larger in young (5026 ± 1613 μm^2^) compared with old (2959 ± 1095 μm^2^; *P* < 0.001) and ∼37% larger in males (4622 ± 1515 μm^2^) compared with females (3363 ± 1709 μm^2^; *P* = 0.003) (Fig. 2A). There was also a clear leftward shift in the size distribution of the MyHC II fibers in older adults, but no age-related shift was observed in the MyHC I fibers (Fig. 2B).

**Fig. 2.**
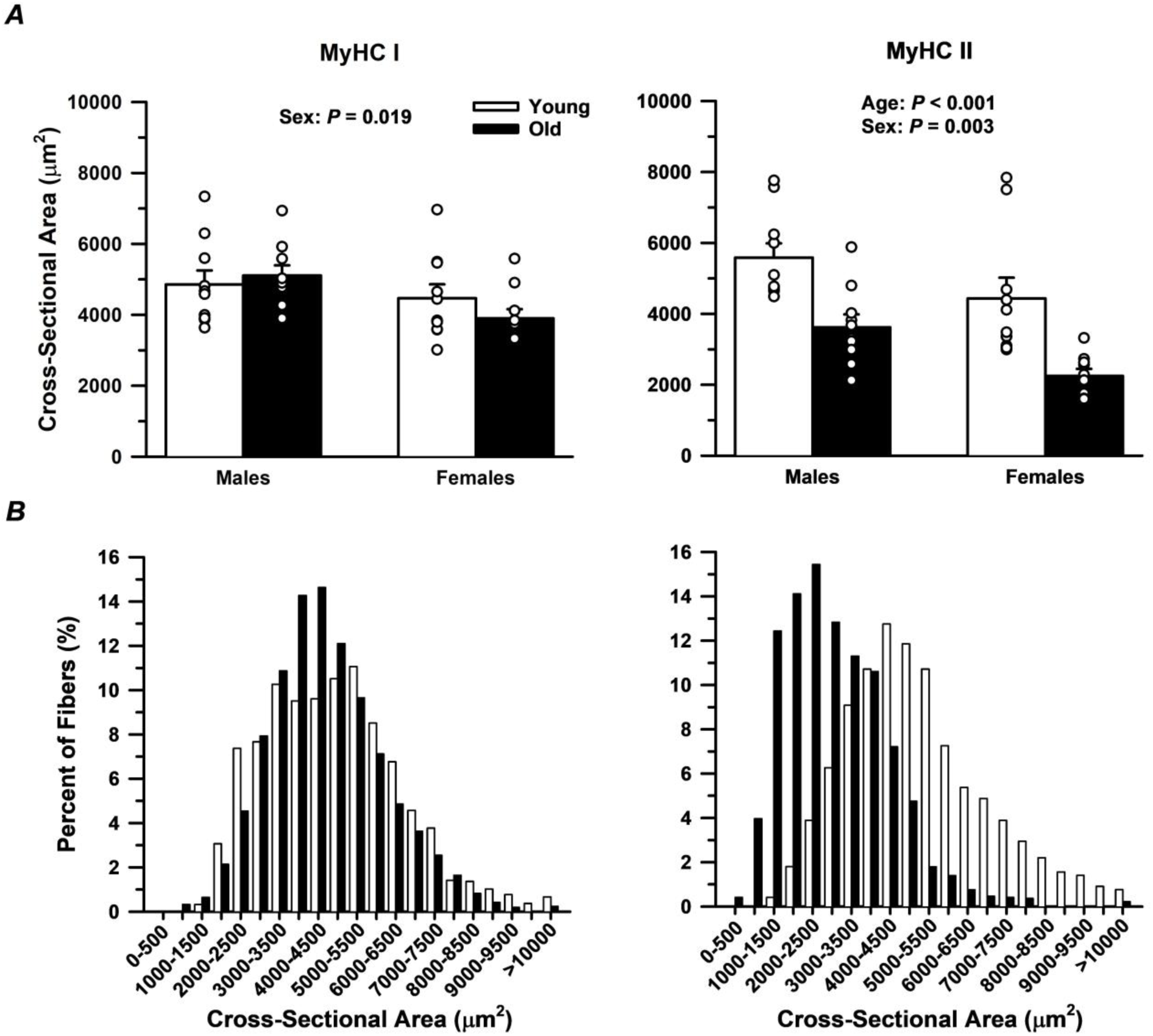
Cross-sectional area (CSA) of slow MyHC I and fast MyHC II fibers. *A*: Males had larger MyHC I fibers compared with females, but there were no age differences. MyHC II fibers were larger in males compared with females, and smaller in older compared with young adults. *B*: A size cluster analysis including a total of 4218 MyHC I fibers and 4055 MyHC II fibers revealed a leftward shift in fast fiber size with ageing, whereas no shift was observed in the slow fibers. Significance level *P* < 0.05. Each dot represents data from an individual participant. Values are displayed as means ± SEM.

### Relative abundance of myonuclei expressing markers of cellular senescence

The percentage of γH2AX^+^/HMGB1^−^ myonuclei in the MyHC I fibers did not differ between young (5.2 ± 2.1%) and old (4.9 ± 3.2%; *P* = 0.765), but females (6.1 ± 2.8%) exhibited a greater percentage of γH2AX^+^/HMGB1^−^ myonuclei compared with males (4.1 ± 2.0%; *P* = 0.015) (Fig. 3A). The percentage of γH2AX^+^/HMGB1^−^ myonuclei in the MyHC II fibers did not differ between young (4.6 ± 2.5%) and old (4.6 ± 2.4%; *P* = 0.967) or between males (4.4 ± 2.4%) and females (4.8 ± 2.5%; *P* = 0.539) (Fig. 3A). There was a significant age by sex interaction in the percentage of γH2AX^+^/HMGB1^−^ myonuclei (*P* = 0.041); however, pairwise comparisons did not reveal any age differences within each sex, nor any sex differences within each age group. When the abundance of γH2AX^+^/HMGB1^−^ myonuclei was expressed per fiber, there were no differences in the MyHC I fibers between young (0.11 ± 0.04) and old (0.10 ± 0.05; *P* = 0.570), or between males (0.10 ± 0.04) and females (0.11 ± 0.04; *P* = 0.322) (Fig. 3B). Similarly, no differences in γH2AX^+^/HMGB1^−^ myonuclei per fiber were observed in the MyHC II fibers between young (0.11 ± 0.06) and old (0.09 ± 0.04; *P* = 0.084), or between males (0.11 ± 0.06) and females (0.09 ± 0.04; *P* = 0.294) (Fig. 3B).

**Fig. 3.**
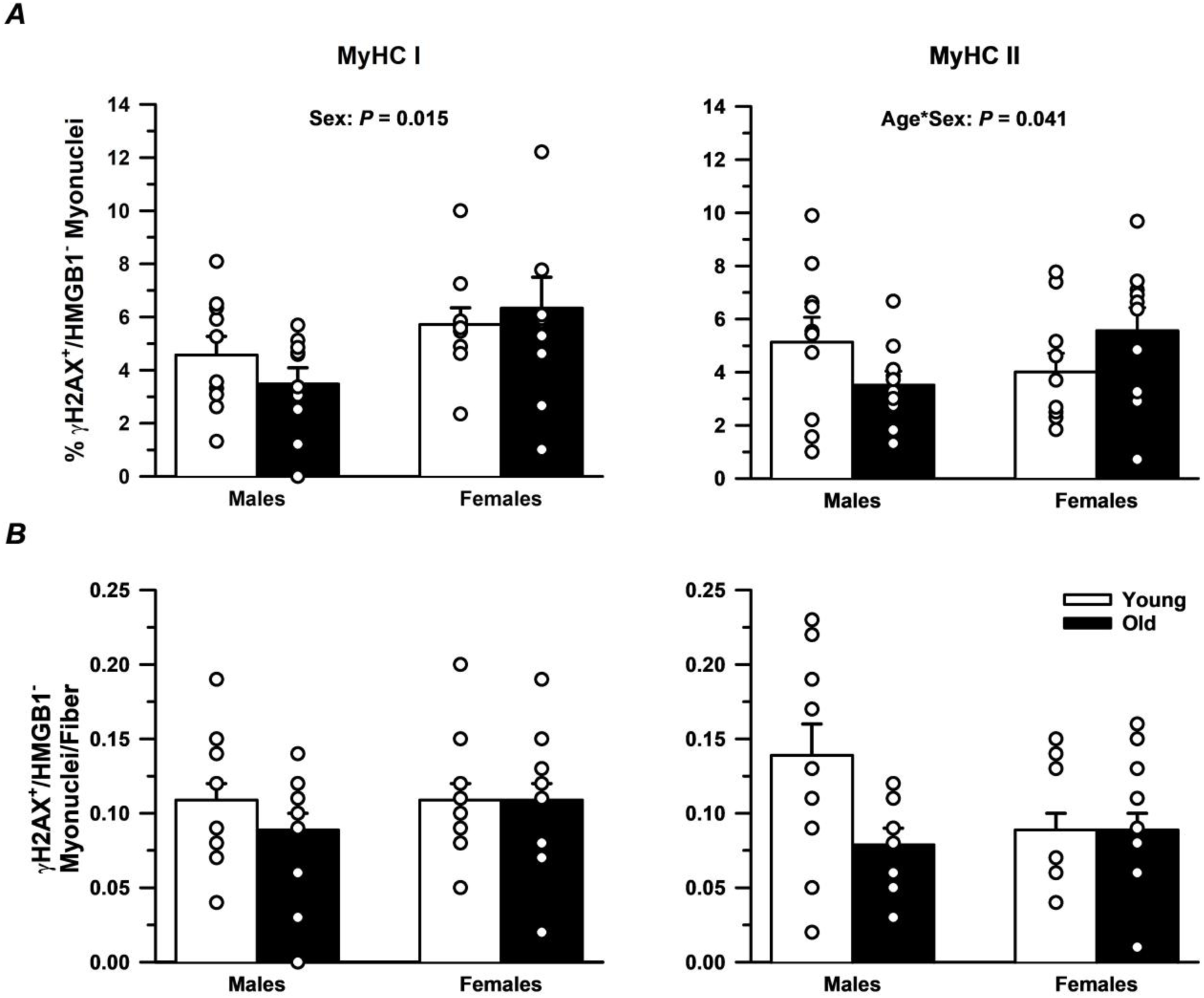
Relative abundance of γH2AX^+^/HMGB1^−^ myonuclei in slow MyHC I and fast MyHC II fibers. *A*: There were no age differences in the percentage of γH2AX^+^/HMGB1^−^ myonuclei in the MyHC I or MyHC II fibers. However, females exhibited a higher percentage of γH2AX^+^/HMGB1^−^myonuclei in the MyHC I fibers compared with males. *B*: There were no age or sex differences in either the MyHC I or MyHC II fibers when the abundance of γH2AX^+^/HMGB1^−^ myonuclei was expressed per fiber. Significance level *P* < 0.05. Each dot represents data from an individual participant. Values are displayed as means ± SEM.

Data for the abundance of myonuclei expressing each senescence marker individually are presented in Table 2. The percentage of γH2AX^+^ myonuclei for the MyHC I fibers was greater in the old (86.1 ± 6.7%) compared with young (80.4 ± 8.0%; *P* = 0.021) but did not differ between males (82.6 ± 7.8%) and females (83.9 ± 8.0%; *P* = 0.615) (Table 2). Similarly, the percentage of γH2AX^+^ myonuclei for the MyHC II fibers was greater in the old (90.6 ± 5.6%) compared with young (83.9 ± 9.9%; *P* = 0.014) but did not differ between males (87.2 ± 7.7%) and females (87.3 ± 9.7%; *P* = 0.987) (Table 2). In contrast, the percentage of HMGB1^−^ myonuclei for the MyHC I fibers did not differ between young (9.9 ± 5.0%) and old (8.8 ± 4.8%; *P* = 0.440) but females (11.1 ± 4.5%) had a greater abundance compared with males (7.7 ± 4.7%; *P* = 0.029) (Table 2). The percentage of HMGB1^−^ myonuclei for the MyHC II fibers did not differ between young (8.9 ± 3.5) and old (7.2 ± 3.5; *P* = 0.109) or between males (7.0 ± 3.1%) and females (9.1 ± 3.7%; *P* = 0.052) (Table 2).

**Table 2:**
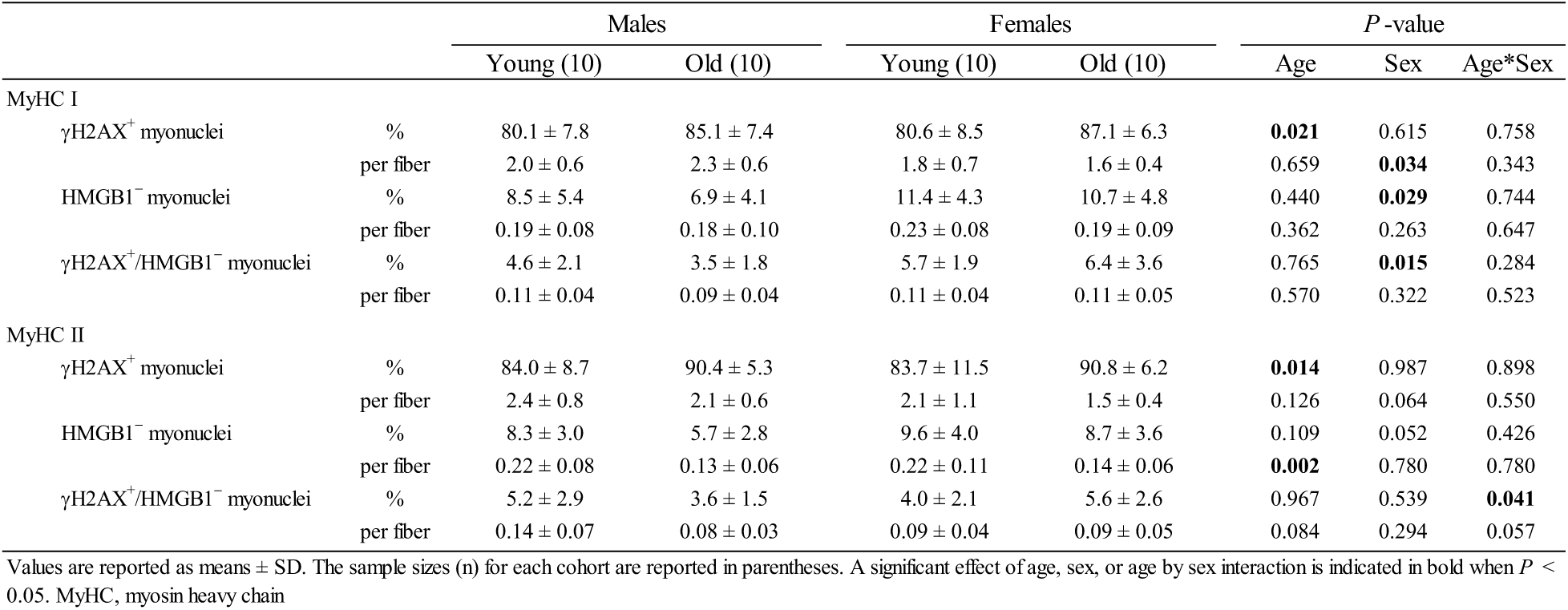
Prevalence of myonuclei expressing markers of cellular senescence.

### Single-nucleus RNA-sequencing (snRNA-seq) data from young and older adult skeletal muscle

To provide a deeper understanding of the prevalence of senescent myonuclei and complement the histological analyses, we re-analyzed a publicly available snRNA-seq dataset (GSE167186). Consistent with the histology findings, *HMGB1* expression did not differ between young and old myonuclei, nor across myonuclei enriched for different myosin types (Figure 4A). Additional genes, including *p21/Cdkn1a* (Fig. 4B) and *p16/Cdkn2a* (Fig. 4C), were evaluated as these are commonly used markers of senescence. Expression of *p16/Cdkn2a* was essentially non-existent in myonuclei regardless of age and fiber type (Figure 4C). Consistent with Perez et al. (2022), a slightly greater proportion of *p21/Cdkn1a*-enriched myonuclei were found in the older adults (MyHC I: young = 0.06%, old = 1.48%; MyHC II: young = 0.27%, old = 3.15%) (Figure 4B). Because *p21* upregulation in murine muscle fibers has been shown to coincide with induction of senescence-associated secretory phenotype (SASP) components and a broader senescence program (Englund et al., 2023), we next evaluated whether the age-related emergence of *p21*-expressing myonuclei was accompanied by a broader senescence signature across myonuclei irrespective of fiber type. Recognizing that snRNA-seq can have limited sensitivity for low-abundance transcripts, we curated a list of SASP- and senescence-related genes based on non-negligible expression in the dataset. Overall, SASP-related gene expression did not differ with age; however, most genes in the curated list were modestly upregulated in myonuclei from young compared with older adults (Fig. 4D).

**Fig. 4.**
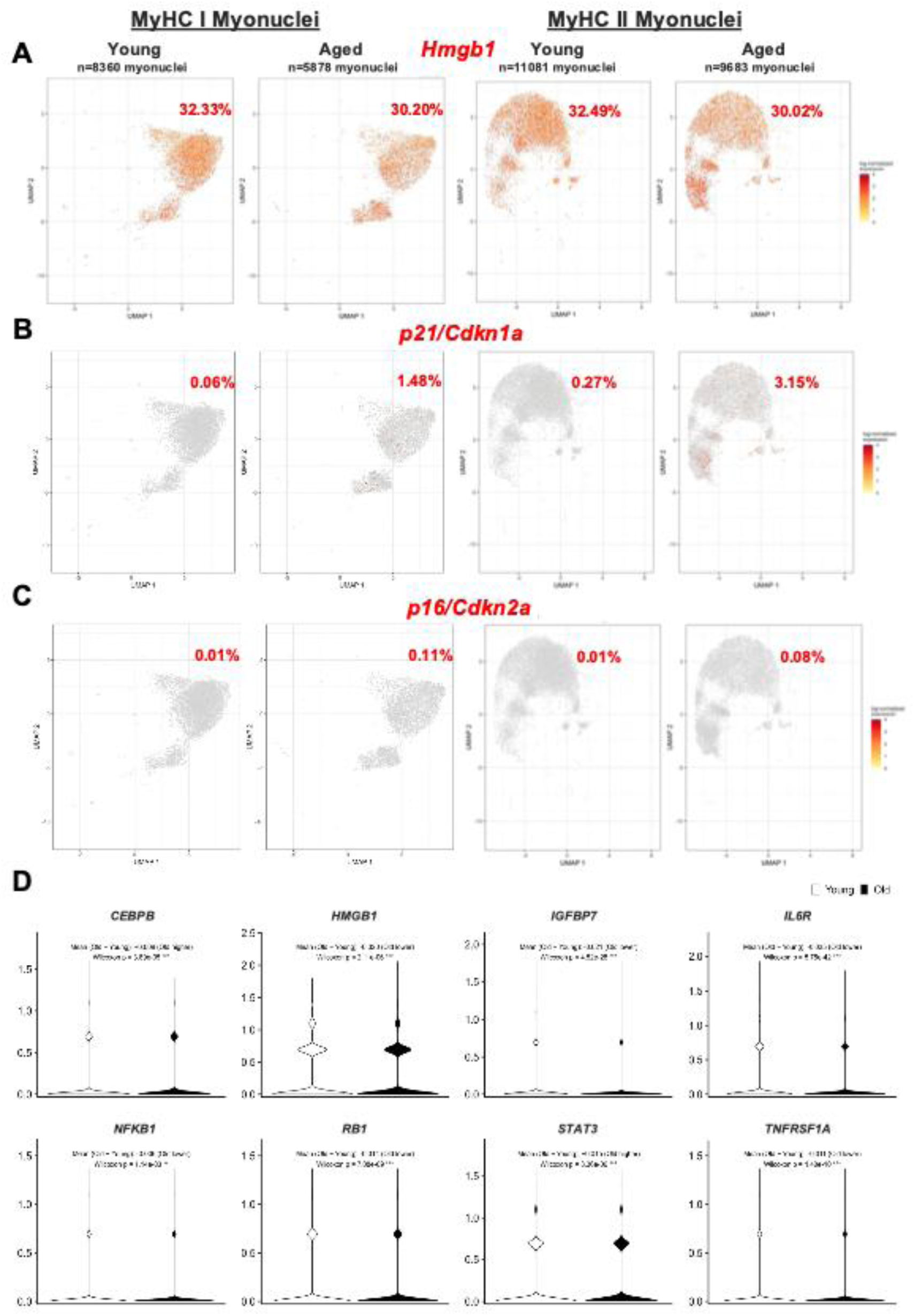
Analysis of senescence-associated markers in single nucleus RNA-sequencing data from Perez et al. (2022) using *vastus lateralis* samples of young and older males. *A*: UMAPs showed *HMGB1* expression did not differ with age or between myonuclei from associated MyHC fiber types. *B*: UMAPs showed *p21/Cdkn1a* expression was higher in the old compared with young in both fiber types. *C*: UMAPs showed *p16/Cdkn2a* expression did not differ with age or between myonuclei from associated MyHC fiber types. *D*: Violin plots representing the expression of senescence program markers myonuclei according to age.

### Myonuclear content

Myonuclei per fiber in MyHC I fibers did not differ between young (2.3 ± 0.7) and old (2.3 ± 0.6; *P* = 0.828), but males (2.6 ± 0.6) had 30% more myonuclei per fiber than females (2.0 ± 0.6; *P* = 0.013) (Fig. 5A). In MyHC II fibers, the young (2.6 ± 1.0) had 30% more myonuclei per fiber compared with old (2.0 ± 0.6; *P* = 0.022), with no difference between males (2.6 ± 0.7) and females (2.1 ± 1.0; *P* = 0.052) (Fig. 5A). A size-cluster analysis performed in both the MyHC I (n = 4218) and MyHC II (n = 4055) fibers revealed no apparent age differences in myonuclear content across the range of fiber sizes (Fig. 5B). Consistent with this, the slope of the linear relationship between myonuclear content and fiber size did not differ between young and old in either fiber type (MyHC I: *P* = 0.889; MyHC II: *P* = 0.547). However, exclusively in muscle from older adults, we observed a distinct group of MyHC II fibers with CSA of 0–1,000 µm² (∼4% of all fibers) that exhibited the lowest myonuclear content across all fiber sizes.

**Fig. 5.**
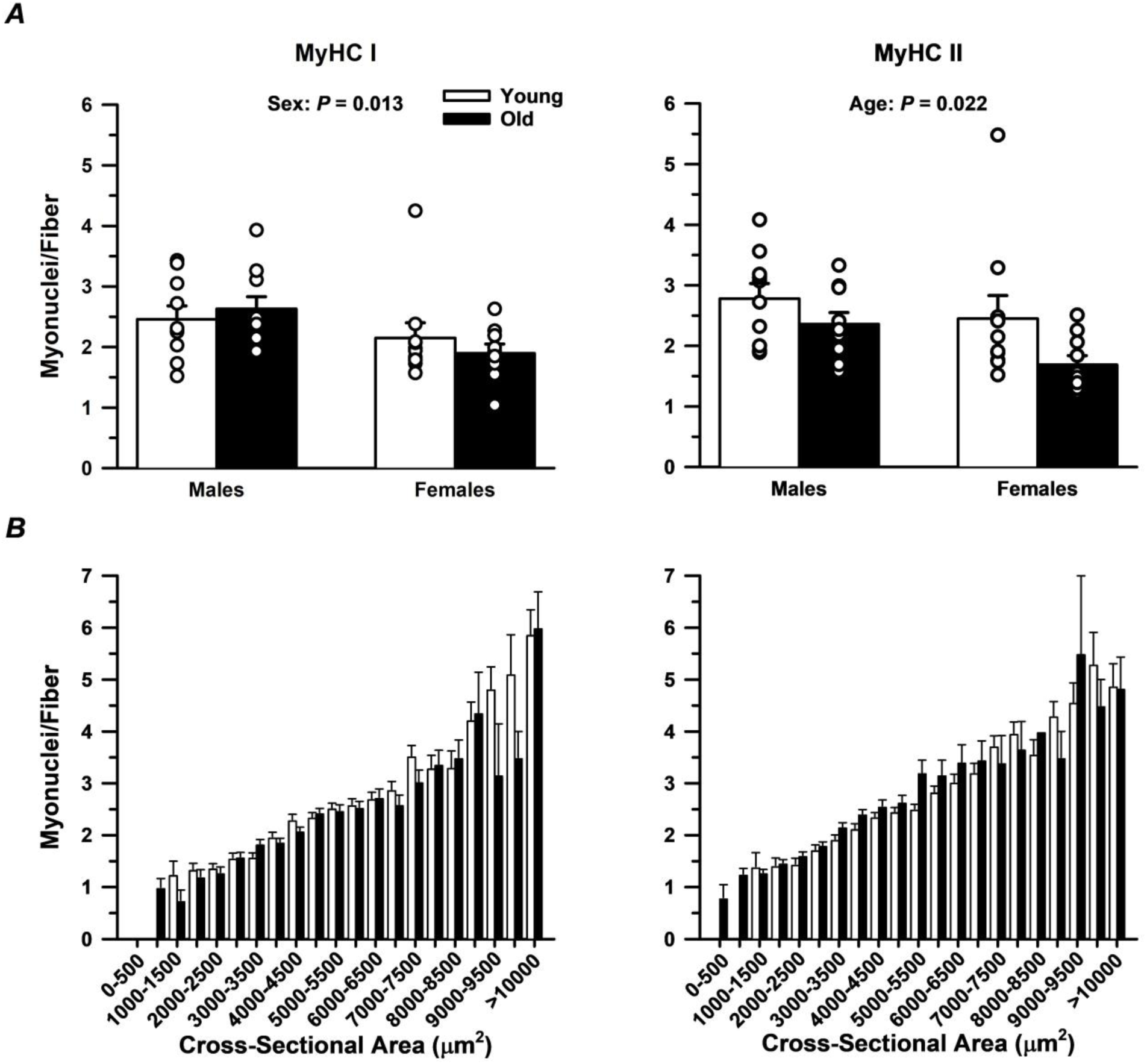
Myonuclear content in slow MyHC I and fast MyHC II fibers. *A*: There was no difference in myonuclei per fiber in the MyHC I fibers between young and old, but males had a greater number of myonuclei per fiber compared with females. In MyHC II fibers, young had a greater number of myonuclei per fiber compared with old, but there were no sex differences. *B*: A size cluster analysis including a total of 4218 MyHC I fibers and 4055 MyHC II fibers showed no apparent age difference in myonuclear content among fibers of similar size within each fiber type. Significance level *P* < 0.05. Each dot represents data from an individual participant. Values are displayed as means ± SEM.

### Myonuclear domain

Myonuclear domain in MyHC I fibers did not differ between young (2191 ± 422 μm^2^) and old (2160 ± 341 μm^2^; *P* = 0.805) or between males (2250 ± 413 μm^2^) and females (2101 ± 336 μm^2^; *P* = 0.227) (Fig. 6A). In contrast, myonuclear domain in MyHC II fibers was ∼53% larger in the young (2201 ± 454 μm^2^) compared with old (1440 ± 355 μm^2^; *P* < 0.001), and ∼24% larger in males (2016 ± 492 μm^2^) compared with females (1625 ± 561 μm^2^; *P* < 0.001) (Fig. 6A). A size-cluster analysis revealed no apparent age-related differences in myonuclear domain across the range of fiber sizes studied in either MyHC I or MyHC II fibers (Fig. 6B). Consistent with this, the slope of the linear relationship between myonuclear domain and fiber size did not differ between young and old in either fiber type (MyHC I: *P* = 0.140; MyHC II: *P* = 0.873).

**Fig. 6.**
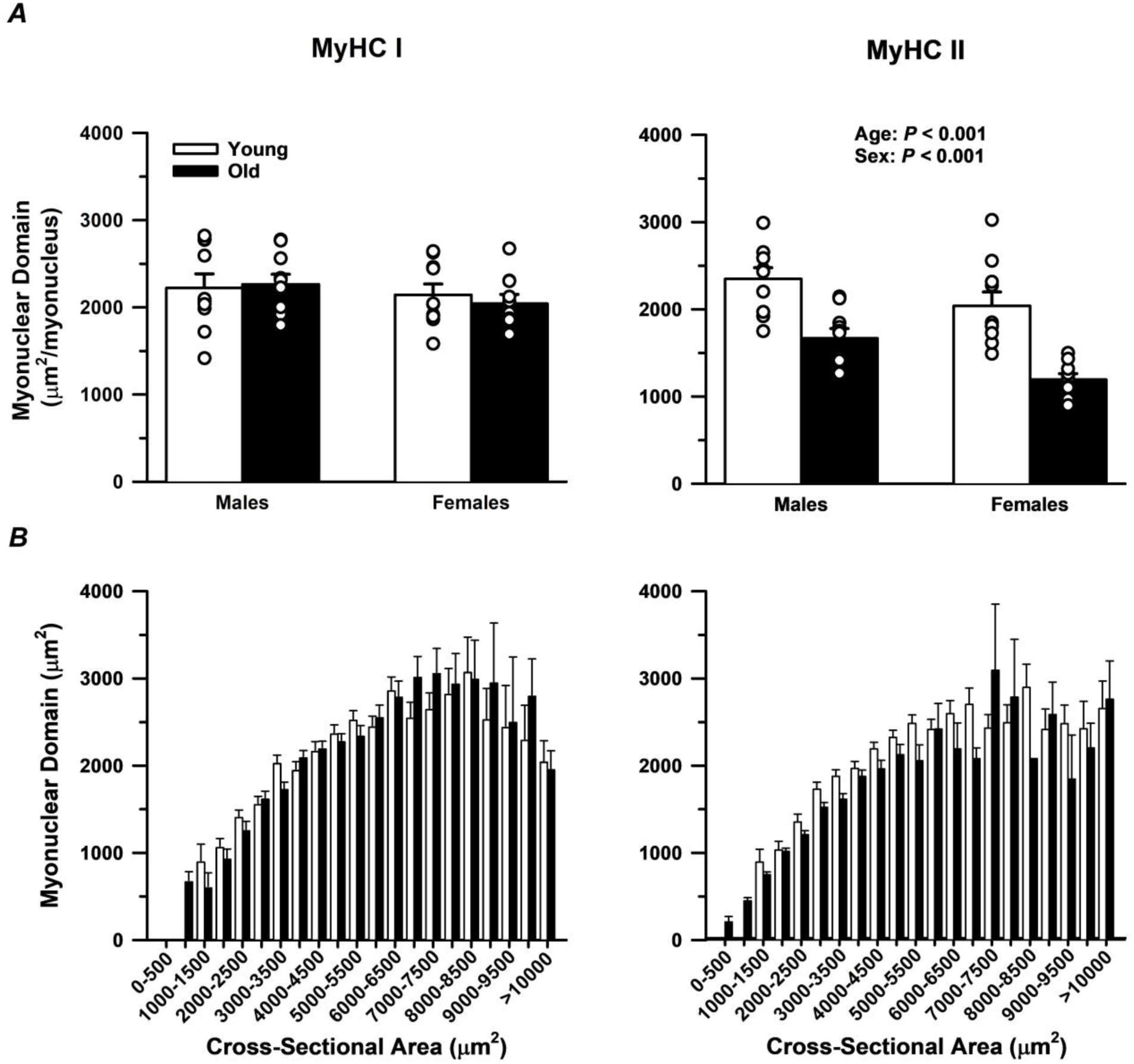
Myonuclear domain in slow MyHC I and fast MyHC II fibers. *A*: There were no age or sex differences in myonuclear domain of the MyHC I fibers. In contrast, myonuclear domain of the MyHC II fibers was larger in young compared with old, and larger in males compared with females. A size cluster analysis including a total of 4218 MyHC I fibers and 4055 MyHC II fibers showed no apparent age difference in myonuclear domain among fibers of similar size within each fiber type. The histograms further illustrate the considerable flexibility in myonuclear across fiber sizes for both fiber types. Significance level *P* < 0.05. Each dot represents data from an individual participant. Values are displayed as means ± SEM.

### Correlations between abundance of myonuclei expressing markers of cellular senescence and myonuclear content with fiber CSA

All subjects were grouped together to examine the relationship between myonuclei expressing markers of cellular senescence, myonuclear content, and myonuclear domain with fiber CSA. No correlation was observed for either the MyHC I (*P* = 0.095) or MyHC II (*P* = 0.367) fibers between the percentage of γH2AX^+^/HMGB1^−^ myonuclei and fiber CSA (Fig. 7A). Although the percentage of γH2AX^+^ myonuclei was greater in old compared with young in both fiber types (Table 2), there were no associations between the percentage of γH2AX^+^ myonuclei and fiber CSA for either the MyHC I (R^2^ = 4 x10^-6^; *P* = 0.99) or MyHC II (R^2^ = 0.05; *P* = 0.151) fibers. In contrast, myonuclear content and myonuclear domain of both the MyHC I (*P* < 0.001) and MyHC II (*P* < 0.001) fibers were significantly correlated with fiber CSA (Fig. 7B-C).

**Fig. 7.**
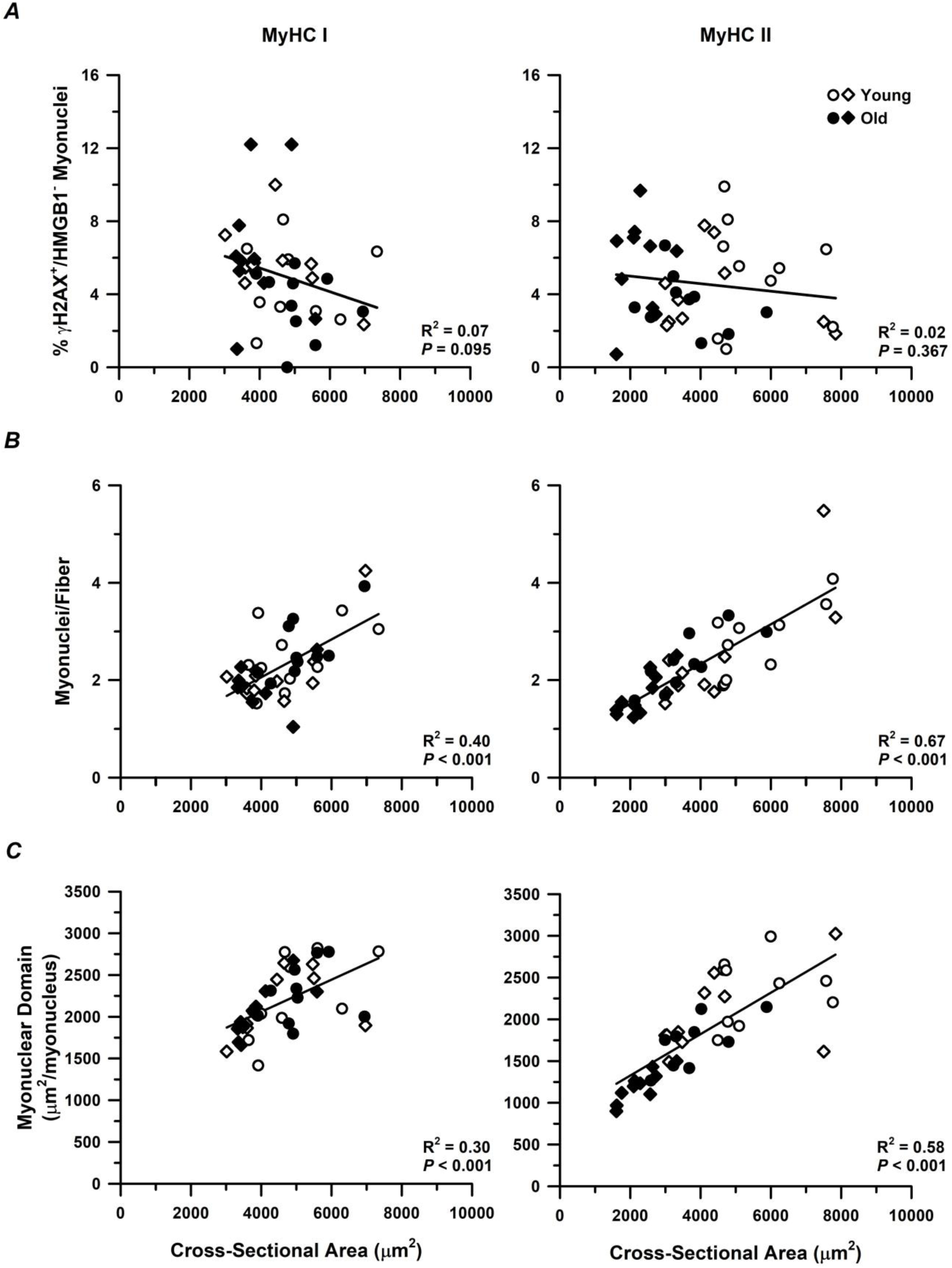
Associations between the prevalence of γH2AX^+^/HMGB1^−^ myonuclei, myonuclear content, and myonuclear domain with fiber CSA. *A*: No associations were observed between the prevalence of γH2AX^+^/HMGB1^−^ myonuclei and fiber CSA in either fiber type. *B-C*: Myonuclear content and myonuclear domain showed moderate-to-strong associations with fiber CSA in both fiber types. Young and old males and females were grouped together for these correlation analyses, and each symbol represents data from an individual participant. Circles (○) represent data from males and diamonds (◇) represent data from females. Significance level *P* < 0.05.

## DISCUSSION

This study aimed to determine whether there is an increased abundance of senescent myonuclei in skeletal muscle fibers of older compared with young adults using a multi-senescence marker approach. As expected, older adults exhibited hallmark features of aging skeletal muscle, including lower thigh lean mass (Table 1) and marked atrophy of fast MyHC II fibers (Fig. 2B). Contrary to our hypothesis, the prevalence of γH2AX^+^/HMGB1^-^ myonuclei did not differ between young and older adults in either slow MyHC I or fast MyHC II fibers and was not associated with CSA in either fiber type (Fig. 3 and Fig. 7A). When the senescence markers were analyzed individually, the prevalence of γH2AX^+^ myonuclei was greater in older compared with young adults in both fiber types (Table 2); however, there was no association of these markers with fiber CSA. In contrast, myonuclear content was lower in fast MyHC II fibers of older compared with young adults and was strongly associated with CSA in both fiber types (Fig. 5A and Fig. 7B). These data suggest that age-related fiber type-specific atrophy cannot be attributed to an increased accumulation of senescent myonuclei but instead implicate myonuclear loss as an important feature of fiber type-specific atrophy with aging.

### Abundance of senescent myonuclei is unable to explain fiber type-specific atrophy with aging

As expected, fast MyHC II fiber size in the older adults was markedly smaller than young adults (Fig. 2A) which was accompanied by a leftward shift in the fiber size distribution (Fig. 2B). In contrast, no age differences were observed in the slow MyHC I fibers (Fig. 2A-B). These findings are consistent with a large body of literature suggesting that fast fibers are more susceptible to reductions in size with aging in both males and females (Coggan et al., 1992; Horwath et al., 2024; Karlsen et al., 2019; Klitgaard et al., 1990; Kramer et al., 2017; Larsson et al., 1978; Lee et al., 2024; Lexell et al., 1988; Nilwik et al., 2013; Soendenbroe et al., 2020). Although the mechanisms contributing to fast fiber atrophy with aging are not fully understood, the accumulation of senescent markers within skeletal muscle has been suggested to contribute to age-related losses in muscle mass and physical function (Fielding et al., 2022; Zhang et al., 2022).

The prevalence of γH2AX^+^/HMGB1^−^ myonuclei did not differ between young and older adults in either slow MyHC I or fast MyHC II fibers (Fig. 3A, B). However, when the senescence markers were analyzed separately, the slow and fast fibers from older adults exhibited a greater relative abundance of γH2AX^+^ myonuclei compared to young (Table 2). Notably, the ∼80-91% γH2AX^+^ myonuclei observed (Table 2) is considerably higher than the ∼34-36% reported by Dungan et al. (2020), but comparable to what was reported in skeletal muscle-derived fibroadipogenic progenitors from young and old mice (Zhang et al., 2022). The potential explanation for this discrepancy is unclear; however, the immunohistochemical approach used in the present study and by Dungan et al. (2020) only permits the binary assessment of the presence or absence of γH2AX. The number of γH2AX foci has been used to quantify the extent of DNA damage (Mah et al., 2010), but foci number cannot be determined in tissue cross-sections due to the limited view of each nucleus. While the prevalence of γH2AX^+^ myonuclei in the present study is higher, the severity of DNA damage within individual nuclei may differ with age and fiber type and warrants further investigation.

Dungan et al. (2020) performed a comprehensive analysis to identify the presence of multiple senescence markers in muscle tissue cross-sections from young and older adults and found no age difference in the prevalence of myonuclei positive for γH2AX^+^. The findings from Dungan et al. (2020) are in contrast to the present study and other studies that found an age-related increase in the prevalence of HMGB1^−^ nuclei or telomere-associated foci in aged muscle tissue (da Silva et al., 2019; Zhang et al., 2022). The reasons for the disparate findings between studies are unclear, but it should be noted that neither da Silva et al. (2019) nor Zhang et al. (2022) used a marker to identify fiber borders to delineate between myonuclei and the nuclei of other mononuclear cells residing in skeletal muscle. Mononuclear cells comprise ∼30-40% of all nuclei within skeletal muscle tissue samples (Dos Santos et al., 2020; Von Walden et al., 2020). Thus, not using a muscle morphology marker makes it difficult to determine whether the age-related reduction in muscle fiber size observed in these previous studies was due to an increased prevalence of myonuclei expressing markers of senescence, or other mononuclear cells residing in skeletal muscle. Due to the larger relative abundance of myonuclei in skeletal muscle, an increased prevalence of myonuclei expressing senescence markers in the fibers of older adults can likely have more profound implications on muscle mass and function compared to other senescent interstitial cells. Although we did not observe age differences in the prevalence of γH2AX^+^/HMGB1^−^ myonuclei in either the slow or fast fibers, it is important to assess fiber type differences due to the susceptibility of the fast fibers to atrophy and reduced contractile function with aging.

Although no age differences were observed in either fiber type for an increased prevalence of senescent myonuclei, we were still interested in investigating whether there was a relationship between fiber size and senescent myonuclei. In the present study, there was no relationship between the prevalence of γH2AX^+^/HMGB1^−^ myonuclei and fiber CSA for either fiber type (Fig. 6A). This is in contrast to the findings of da Silva et al. (2019) where a significant association was found between the prevalence of HMGB1^−^ nuclei and muscle fiber diameter in older mice. A potential reason for the conflicting findings includes, as mentioned previously, not using a muscle morphology marker to distinguish myonuclei from other muscle-resident mononuclear cells. The conflicting findings could also be due to a species difference in which senescent myonuclei may play a larger role in regulating muscle fiber size in rodents compared to humans; however, this explanation seems unlikely given recent evidence suggesting that the senescence phenotype in skeletal muscle is conserved between preclinical models and humans (Zhang et al., 2022).

A limitation of the present study is that only two senescence-associated nuclear markers, γH2AX and HMGB1, were used to assess age differences in the prevalence of senescent myonuclei. These markers were chosen because they reflect DNA damage and chromatin instability, respectively, which could compromise myonuclear function and contribute to reduced fiber size with aging. The use of multiple markers to identify senescent cells has been proposed because many widely used markers, including senescence-associated β-galactosidase (SA β-Gal), p16, and p21, can be pleiotropic (Gonzalez-Gualda et al., 2021; Gorgoulis et al., 2019; Ogrodnik et al., 2024). Our re-analysis of the human muscle snRNA-seq data from Perez et al. (2022) reinforces an induction of *p21* with age in myonuclei enriched for MyHC II genes, and to a lesser extent in MyHC I-expressing myonuclei (Fig. 4B). However, the low prevalence of other senescence-associated transcripts across myonuclei in older muscle (Fig. 4D) suggests that *p21* may play a role in aging skeletal muscle dysfunction, such as fibrosis and/or mitochondrial dysfunction (Englund et al., 2023), that could arise independent from senescence. A key feature of senescent cells is their resistance to clearance from tissues. If this principle applies to nuclei within a syncytium, then the apparent lack of overt senescence-associated features in myonuclei may be permissive for myonuclear removal during fiber atrophy across the lifespan.

The present study also aimed to explore potential sex differences in the prevalence of senescent myonuclei in the slow and fast fibers. Interestingly, females exhibited a higher prevalence of γH2AX^+^/ HMGB1^−^ myonuclei in the slow fibers compared to males irrespective of age, with no sex differences in the fast fibers (Fig. 3A). Given that there was no sex difference in the prevalence of γH2AX^+^ myonuclei in the slow fibers, this finding may be explained, in part, by the higher relative abundance of HMGB1^−^ myonuclei in the slow fibers of females compared to males (Table 2). These results align with previous studies investigating processes associated with cellular senescence in other cell types, which demonstrated that females had reduced capacity for DNA damage repair in peripheral blood mononuclear cells (PBMCs) (Trzeciak et al., 2008) and lymphocytes (Rall-Scharpf et al., 2021). Taken together, these findings suggest that a lower capacity for DNA damage repair may lead to an increased accumulation of DNA damage which would manifest as an increased prevalence of senescent myonuclei in females compared to males. However, this is in contrast to other studies in human PBMCs (Garm et al., 2013) and lymphocytes (Sharma et al., 2015) reporting that biological sex does not influence DNA damage repair capacity. Furthermore, a recent study exploring associations between senescence markers with muscle morphology and muscle function across the lifespan found a moderate negative relationship (*r* = - 0.44) between HMGB1^+^ nuclei and age in females, but not males, suggesting a sex-specific loss in nuclear HMGB1 expression with aging (Habiballa et al., 2024). Habiballa et al. (2024) also found HMGB1 was positively associated with grip strength in females (*r* = 0.52), but not males. The translocation of HMGB1 from the nucleus to the extracellular space is known to promote inflammatory responses (Yamada & Maruyama, 2007). Thus, the loss in HMGB1 with advancing age in females may lead to increased inflammation which has been shown to be associated with muscle atrophy (Schaap et al., 2006) and reduced muscle function (Ferrucci et al., 2002). Given the conflicting findings on sex-specific expression of senescent markers, and the unclear reason for the increased prevalence of senescent myonuclei in the slow fibers of females irrespective of age, our data highlight the need for further research focused on expanded fiber type-specific senescence marker expression and its contribution to age-related declines in muscle mass and function in females compared with males.

### Myonuclei are lost in the fast fibers of older adults and are associated with a smaller fiber size

We found lower average myonuclear content in the fast fibers from old compared with young adults, with no age-related difference observed in slow fibers (Fig. 4A). The lower myonuclear content in fast fibers was accompanied by marked reductions in fiber CSA and myonuclear domain. As expected, correlation analyses revealed moderate-to-strong associations between fiber CSA and myonuclear content in both fibers types when young and older males and females were evaluated together (Fig. 6B) (Horwath, Envall, et al., 2021; Horwath, Moberg, et al., 2021; Karlsen et al., 2015; Verdijk et al., 2010). Similarly, fiber CSA was moderately-to-strongly associated with myonuclear domain (Fig. 6C). To determine whether the lower myonuclear content and smaller myonuclear domain in MyHC II fibers from older adults were attributed solely to their smaller fiber size (Karlsen et al., 2015), we performed a size-cluster analysis across all 4,022 fibers. This analysis indicated that myonuclear content did not appear to differ appreciably with age across the full range of similarly sized fibers, and that myonuclear content scaled similarly with fiber CSA in young and older adults. Thus, the lower myonuclear content and smaller myonuclear domain observed in fast fibers from older adults can largely be explained by the markedly greater prevalence of small fast fibers (i.e. age-associated atrophy) (Fig. 2B). Together, these data suggest that fast fiber atrophy corresponds closely with the loss of myonuclei; however, whether the loss of myonuclei precedes fiber atrophy or vice versa cannot be determined definitively. We speculate that reductions in fiber size occur more rapidly than myonuclear removal for two reasons: 1) this pattern has been observed during unloading and inactivity (Jackson et al., 2012; Murach et al., 2018), and 2) qualitative inspection from our cluster analysis suggests that for a given fiber size, MyHC II fibers from older adults tend to have slightly more myonuclei (Fig. 5B) and slightly smaller myonuclear domains (Fig. 6B), consistent with a more rapid atrophy process relative to myonuclear removal during aging. These findings collectively support the idea that the myonuclear domain, particularly in the fast fibers, is more flexible than previously suggested (Bagley et al., 2023; Murach et al., 2018).

An age-related decline in fast fiber myonuclear content from the *vastus lateralis* of humans has been reported (Kramer et al., 2017; Verdijk et al., 2014; Verdijk et al., 2016); however, this is not a universal finding as other studies have shown no change (Cristea et al., 2010; Horwath et al., 2024; Kelly et al., 2018; Montenegro et al., 2024) or even an increase (Cristea et al., 2010; Verdijk et al., 2007) with age. Whether myonuclei are lost with aging and/or muscle fiber atrophy was recently debated (Kirby & Dupont-Versteegden, 2022; Kirby & Dupont-Versteegden, 2022; L. Schwartz & K. Gundersen, 2022; L. M. Schwartz & K. Gundersen, 2022) with potential mechanisms including myonuclear apoptosis (Dupont-Versteegden, 2005) and/or autophagy (Luo et al., 2016), consistent with the removal of mitochondria via mitophagy, another DNA-containing organelle (Yamashita & Kanki, 2017). In the present study, we found an age-related increase in the prevalence of γH2AX^+^ myonuclei, with the highest prevalence in MyHC II fibers. It is therefore enticing to speculate that the accumulation of DNA damage in myonuclei with aging, which does not appear to correspond with a senescent phenotype, may initiate myonuclear removal as fast fibers atrophy. Discrepancies in the literature regarding the occurrence of myonuclear loss with age could be attributed to methodological limitations. For instance, discriminating myonuclei from other mononuclear cells within skeletal muscle can be particularly challenging on muscle tissue cross-sections due to alterations in myonuclear shape that may occur with aging and physical activity (Battey et al., 2023; Bruusgaard et al., 2006; Cristea et al., 2010; Murach et al., 2020). Irrespective of the potential explanation for the discrepancies between studies, our findings are consistent with a recent meta-analysis from human and animal studies demonstrating lower myonuclear content and smaller myonuclear domain in fast fibers of older compared with young adults (Rahmati et al. 2022). Collectively, these findings suggest an important role for myonuclei in regulating muscle fiber size (Hansson et al., 2020) and suggest that interventions, such as resistance training, that promote myonuclear accretion may be critical for preserving muscle fiber size and function with age.

## Conclusions

Older adults exhibited marked atrophy of fast fibers which was accompanied by reductions in myonuclear content and myonuclear domain size. Using a multi-marker senescence approach, we found no age-related differences in the prevalence of senescent myonuclei in either fiber type. However, a sex difference in the prevalence of senescent myonuclei was observed in slow fibers only. Fiber CSA was closely associated with myonuclear content, but not with the prevalence of senescent myonuclei, suggesting an important role for myonuclei in regulating muscle fiber size. We conclude that myonuclear loss, rather than the accumulation of senescent myonuclei, is a key feature of fiber type-specific atrophy with aging.

## DATA AVAILABILITY

The data in the present study are available from the corresponding author upon reasonable request.

## ACKNOWLEDGMENTS

We thank Dr. Carolyn Smith for assisting with some of the muscle biopsies and the research participants for volunteering to make this study possible. Thank you to Dr. Davis Englund for helpful discussions regarding senescence markers in the snRNA-seq data.

## AUTHOR CONTRIBUTIONS

CSZ and CWS conceived and designed the experiments; CSZ, ID, and CWS performed experiments; CSZ, ID, PNA, and RGJ3 analyzed the data; CSZ and CWS interpreted the data; CSZ, RGJ3 and CWS prepared the figures; CSZ and CWS drafted the manuscript; CSZ, CMD, RGJ3, KAM, and CWS edited and revised the manuscript; CSZ, ID, PNA, CMD, RGJ3, KAM, and CWS approved the final version of the manuscript.

---

**Table A1:** Number of fibers included in size cluster analyses.

| Size Cluster | MyHC I |  | MyHC II |  |
| --- | --- | --- | --- | --- |
|  | Young | Old | Young | Old |
| <500 | 0 | 0 | 0 | 10 |
| 500-1000 | 1 | 9 | 0 | 82 |
| 1000-1500 | 8 | 16 | 10 | 254 |
| 1500-2000 | 63 | 49 | 38 | 288 |
| 2000-2500 | 149 | 102 | 80 | 315 |
| 2500-3000 | 155 | 177 | 128 | 262 |
| 3000-3500 | 207 | 242 | 185 | 231 |
| 3500-4000 | 192 | 317 | 218 | 217 |
| 4000-4500 | 194 | 325 | 259 | 148 |
| 4500-5000 | 212 | 269 | 241 | 98 |
| 5000-5500 | 223 | 215 | 218 | 38 |
| 5500-6000 | 172 | 159 | 148 | 30 |
| 6000-6500 | 137 | 109 | 110 | 17 |
| 6500-7000 | 93 | 82 | 100 | 11 |
| 7000-7500 | 77 | 58 | 80 | 10 |
| 7500-8000 | 30 | 38 | 61 | 9 |
| 8000-8500 | 29 | 20 | 46 | 1 |
| 8500-9000 | 22 | 11 | 33 | 2 |
| 9000-9500 | 17 | 6 | 30 | 2 |
| 9500-10000 | 9 | 2 | 20 | 2 |
| >10000 | 15 | 7 | 17 | 6 |
| Total | 2005 | 2213 | 2022 | 2033 |

The number of slow MyHC I and fast MyHC II fibers included within each size bin for young and older adults. The number of fibers were pooled across males and females within each age group.

